# Number of mismatches and length of longest match correlate with alignment score in swalign built-in function in MATLAB

**DOI:** 10.1101/2020.07.12.199661

**Authors:** Wenfa Ng

## Abstract

Understanding how one sequence relates to another at the nucleotide or amino acid level allows the derivation of new knowledge regarding the provenance of particular sequence as well as the determination of consensus sequence motifs that informs biological conservation at the sequence level. To this end, local or multiple sequence alignments tools in bioinformatics have been developed to automatically profile two or more nucleotide or amino acid sequence in search of matches in stretches of nucleotides or amino acid sequence that yield an alignment. While alignment score is a common metric for assessing alignment quality, relative difference between alignment scores does not readily correlate with concrete measures such as number of mismatches and length of longest match in alignment. Thus, using swalign local sequence alignment function in MATLAB on 200 alignments between RNA-seq sequence read and reference *Escherichia coli* K-12 MG1655 genome sequence in the sense and antisense direction, this work sought to shed some light on how alignment score from swalign correlates with number of mismatches and length of longest match. Results revealed that number of mismatches negatively correlate with alignment score; thereby, validating theoretical predictions that larger number of mismatches would result in a poorer alignment and lower alignment score. However, dependence of alignment score on other factors such as length of longest match and gap penalty from opening an alignment gap prevents linear relationship to be obtained between number of mismatches and alignment score. On the other hand, length of longest match was found to positively correlate with alignment score as predicted from theoretical understanding. But, data obtained revealed that clusters of data points gather at two regions of the scatter plot involving short matches and low alignment score, as well as long matches and high alignment score. Such clustering and sparseness of data points between the two clusters preclude the elucidation of a linear quantitative relationship between length of longest match and alignment score. Overall, dependence of alignment score of swalign on number of mismatches and length of longest match in alignment match theoretical predictions; thereby, validating the utility of alignment score in indicating the qualitative quality of alignment. However, given that alignment score inherently depends on a multitude of factors, users could not easily discern the quantitative difference in mismatches and length of longest match from relative differences between two alignment scores. Such problems are unlikely to be resolved given the near impossibility of obtaining quantitative linear relationship correlating either number of mismatches or length of longest match with alignment score of a sequence alignment tool.

**Highlights:**

1. Number of mismatches in alignment negatively correlates with alignment score.
2. Length of longest match positively correlates with alignment score.
3. Quantitative linear relationship could not be obtained for alignment score with either number of mismatches or length of longest match.
4. Results validate that swalign tool in MATLAB could quantitatively detect differences in alignment quality and expressed it using alignment score.
5. But, relative alignment score of two alignments remains a nebulous concept with regards to differences in number of mismatches and length of longest match.

## Introduction

Sequence alignment is a foundational tool in bioinformatics for understanding provenance of different genes and determining consensus sequence of particular motifs such as the -35 and -10 box of promoters. The method is also used in determining how sequence reads from transcriptome sequencing matches to their corresponding genes in the reference genome. To help in this endeavour, multiple sequence alignment tools using different algorithms have been published. Typically, sequence alignment tools would present the result of the alignment in both graphical format and an alignment score. However, alignment score remains a nebulous concept without detailed knowledge of the sequence alignment algorithm. Specifically, how the alignment score relates to the number of mismatches and length of longest match are key questions that determine how an alignment score is to be interpreted.

Theoretically, alignment score should negatively correlate with number of mismatches; thus, a high alignment score should correlate with a sequence alignment with few mismatches. On the other hand, alignment score typically would be higher if the length of the longest match in sequence alignment is long; thereby, yielding a positive correlation between alignment score and length of longest match. Overall, in understanding the significance of an alignment score, several parameters that feed into the calculation of alignment score would need to be understood. Besides number of mismatches and length of longest match in the alignment, penalty for opening a gap in the alignment also affects the alignment score.

Thus, the objective of this work is to understand how number of mismatch and length of longest match affects the alignment score of swalign in-built sequence alignment tool in MATLAB. Specifically, the type of alignment investigated in this work concerns how well a sequence read from an RNA-sequencing experiment matches to the reference genome from which RNA transcripts derive. Thus, sequence alignments encountered in this work are those pertaining to short sequence reads (∼50 bases) to a reference genome. Hence, limits exist in which longer matches (hundreds of base-pairs) could not be used to understand how length of longest match influence alignment score. In addition, there is also a limit on the number of mismatch possible in the alignment, which meant that the data points obtained could not fully represent the full range of alignment score possible. Such limitations represent the extend in which results derived from this work should be understood.

## Materials and methods

Sequence reads from RNA-sequencing data of *Escherichia coli* was aligned to the genome sequence of *E. coli* K-12 MG1655 using the swalign built-in function in MATLAB. Swalign is a local sequence alignment tool developed by Smith-Waterman.1 Both the genome sequence and its reverse complement was used in the alignment; thereby, yielding two sequence alignment for each sequence read. The obtained alignment was used in further analysing the number of mismatches and length of longest match in the sequence alignment. Specifically, number of mismatches in the alignment was calculated and length of longest match was determined by a MATLAB programme. A total of 100 sequence reads was analysed in this work.

Number of mismatches was determined by calculating the total number of mismatches in the alignment. On the other hand, length of longest match was calculated by first determining the length of each match within the alignment, and subsequently calculating the match that has the longest length of consecutive nucleotides.

A zip file of the MATLAB programme could be found in the Supplementary information of this preprint.

## Results and Discussion

Analysis of the results revealed that number of mismatches was negatively correlated with alignment score of swalign (Figure 1). Specifically, it could be observed that the higher the number of mismatches, the lower the alignment score. Given that the data points covered the range of mismatches evenly, a negative correlation could be established between alignment score and number of mismatches, which meant that alignment score of swalign function is diagnostic of how well the sequence read align to the genome sequence. The result obtained is also consistent with the theoretical understanding that a greater number of mismatches would result in a lower alignment score. However, even though a negative correlation exists between alignment score and number of mismatches, a direct quantitative relationship between the two variables could not be obtained as alignment score also depends on many other factors, one of which is length of longest match in alignment.

**Figure 1:**
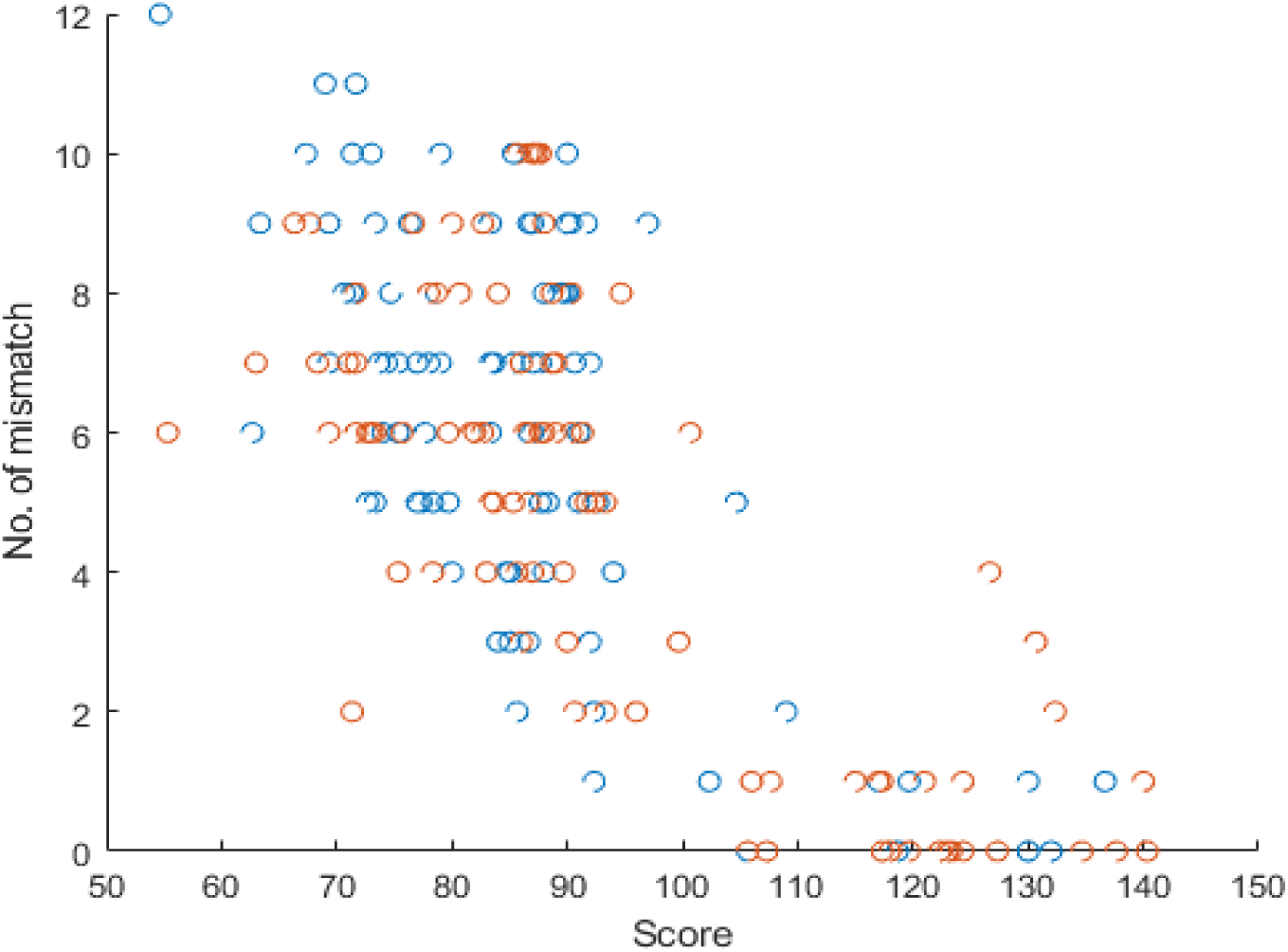
Negative correlation could be seen between number of mismatches in alignment and score of the alignment for sequence reads aligned to a reference genome using swalign function in MATLAB.

For a sequence read of defined length, the greater the number of mismatches, the probability of having a long match would be correspondingly reduced, except for cases where the mismatches are clustered together at a particular section of the sequence read. This is seen in Figure 2 where clusters of data points gather at the short length of match and low alignment score region, as well as the long length of match and high alignment score region. Data available reveals that a positive correlation exists between length of longest match and alignment score, which is consistent with theoretical understanding of how well instances of long matches influence the alignment score.

**Figure 2:**
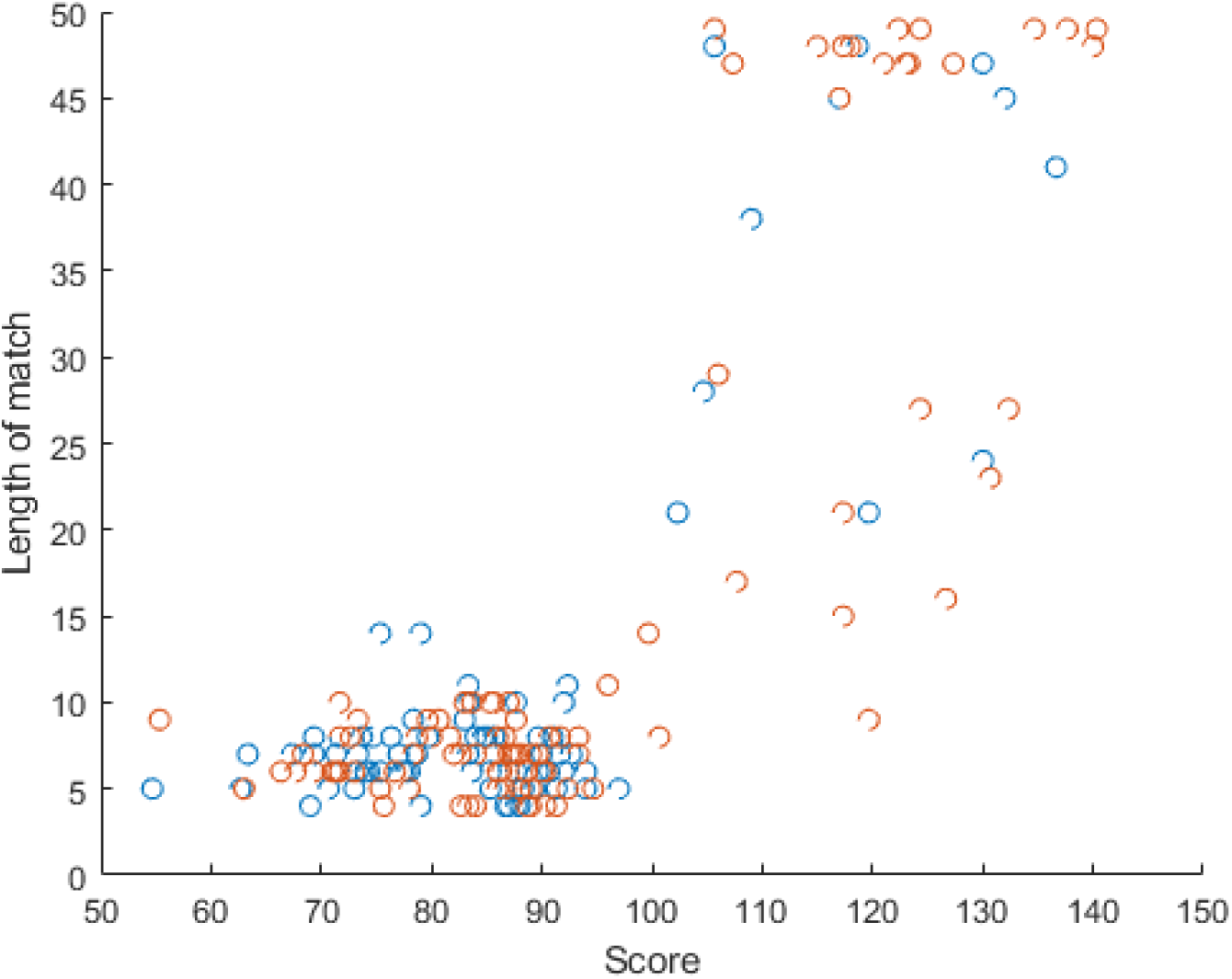
Scatter plot of length of longest match and alignment score for alignment of 100 sequence reads to both the genome sequence of *E. coli* K-12 MG1655 and its reverse complement using swalign function in MATLAB.

However, the data obtained is sparse in the region of intermediate length of longest match and intermediate alignment score, which together with the clustering of data points prevents the elucidation of a quantitative positive correlation between length of longest match and alignment score. More importantly, as mentioned above, alignment score also depends on number of mismatches and the penalty imposed on opening a gap in the alignment; thus, a linear fit could not be obtained between length of longest match and alignment score.

## Conclusions

Overall, the data obtained from aligning 100 sequence reads to the sense and antisense strand of the genome sequence of *E. coli* K-12 MG1655 revealed that number of mismatches in the alignment negatively correlate with alignment score readable from the swalign function which is based on the Smith and Waterman algorithm. However, dependencies of alignment score on other parameters prevent the elucidation of a linear correlation between number of mismatch and alignment score. On the other hand, length of longest match was found to cluster at two regions of the scatter plot relating length of longest match with alignment score, which also precludes the elucidation of a linear relationship between the variables through linear regression. While length of longest match positively correlates with alignment score, dependence of alignment score on other parameters such as gap penalty of opening an alignment gap and number of mismatches also hampers a linear relationship to be obtained between length of longest match and alignment score.

Collectively, elucidation of the dependencies of alignment score on number of mismatches and length of longest match helped verify that swalign function could detect and quantify the relative effect of different number of mismatches and length of longest match on alignment quality. However, given that alignment score also depends on other variables difficult to quantify and the inability to obtain a linear relationship between alignment score and number of mismatches and length of longest match respectively, alignment score remains a nebulous quantity to the end user where its application space is limited to a qualitative understanding of alignment quality of different sequence reads to a reference genome sequence. Finally, while local sequence alignment tools are expected to evolve to better align two sequences with less computational power, utility of alignment score for obtaining a quantitative understanding of quality of alignment remain difficult as the variable is inherently dependent on a multitude of factors.

## Supporting information

MATLAB software supporting the work

## Supplementary information

Source code of the MATLAB programme used for this work is in the programme file that is part of the zip file

