## Supplementary material for "Number of mismatches and length of longest match correlate with alignment score in swalign built-in function in MATLAB": MATLAB software supporting the work: get_longest_match.m

```
function longest_match = get_longest_match(alignment)

    k1 = length(alignment);
    cand =[]; m =1; matchflag = 0; n =1;
    
    for i = 1:k1
        letter = alignment(2,i);
        if contains(letter, '|')
            if matchflag ==0
                start = i;
            end
            matchflag = 1;
            match(n) = letter;
            n = n +1;
        end
        
        if letter ~= '|' && matchflag ==1
            matchflag = 0;
            cand(m).match = match;
            stop = i-1;
            cand(m).start = start;
            cand(m).stop = stop;
            n = 1;
            m = m+1;
            cand(m).matchlength = stop-start+1;
            cand(m).totallength = k1;
            cand(m).distance_from_end = k1-stop;
            clear match
        end
    end
        
        letter = alignment(2, k1);
        
        if letter =='|'
            cand(m).component = match;
            stop = k1;
            cand(m).start = start;
            cand(m).stop = stop;
            cand(m).matchlength = stop-start +1;
            cand(m).totallength = k1;
            cand(m).distance_from_end = k1 - stop;
        end
        
        matchlength_vector = [cand.matchlength];
        [~, index] = max(matchlength_vector);
        result = matchlength_vector(index);
        
        longest_match = result;
            
            
    
    

end
```
