## Supplementary material for "Number of mismatches and length of longest match correlate with alignment score in swalign built-in function in MATLAB": MATLAB software supporting the work: get_mismatches.m

```
function num_of_mismatches = get_mismatches(alignment)

    k1 = length(alignment)
    
    num_of_mismatches = 0;
    
    for i=1:k1
        letter = alignment(2, i);
        if letter == ':'
            num_of_mismatches = num_of_mismatches +1;
        end
    end

end
```
