## Supplementary material for "Number of mismatches and length of longest match correlate with alignment score in swalign built-in function in MATLAB": MATLAB software supporting the work: get_score_and_longest_match.m

```
function get_score_and_longest_match

    load('block_alignment100.mat')
    
    k1 = length(block);
    
    for i =1:100
        Score_forward(i) = block(i).Score_forward;
        Score_reverse(i) = block(i).Score_reverse;
        alignment_forward = block(i).alignment_forward;
        longest_match_forward(i) = get_longest_match(alignment_forward);
        alignment_reverse = block(i).alignment_reverse;
        longest_match_reverse(i) = get_longest_match(alignment_reverse);
    end
    
    scatter(Score_forward, longest_match_forward)
    hold on
    scatter(Score_reverse, longest_match_reverse)
    xlabel('Score'); ylabel('Length of match');
    
    

end
```
