## Supplementary material for "Number of mismatches and length of longest match correlate with alignment score in swalign built-in function in MATLAB": MATLAB software supporting the work: main.m

```
function main

    load('block.mat')
    load('mg1655.mat')
    
    genomeseq = gbfile.Sequence;
    genomeseqrc = seqrcomplement(genomeseq);
    
    k1 = length(block);
    
    for i=1:100
        read = block(i).Sequence;
        read = lower(read);
        [Score_forward, alignment_forward] =swalign(genomeseq, read);
        [Score_reverse, alignment_reverse] =swalign(genomeseqrc, read);
        block(i).Score_forward = Score_forward;
        block(i).alignment_forward = alignment_forward;
        block(i).Score_reverse = Score_reverse;
        block(i).alignment_reverse = alignment_reverse;
    end
   

end
```
